# Computational Assessment on Catalytic Activity of PET Hydrolase

**DOI:** 10.1101/2023.05.30.542930

**Authors:** Igor Nelson, Rafael Ramos

**Affiliations:** University of Coimbra, 200911124; University of Coimbra, 2022129995

**Keywords:** Hydrolase, enzymatic degradation, plastic recycling, protein engineering, polyethylene terephthalate (PET)

## Abstract

**Background:** PET hydrolase from *Ideonella sakaiensis* might provide a response for PET accumulation in the environment. In this project some previously studied mutations were implemented and their performance was evaluated via computational methods with tools such as Modeller, HADDOCK, PyMOL and Gromacs. One possible mutation that could lead to improved catalytic activity was proposed.

**Results:** PET hydrolase DM S209F W130H and I179 provide interesting binding results with studied ligands, however a solution that combines both mutations does not seem viable, since the binding cleft becomes occluded. Following the same rationale, the triple mutant S209F W130H I179Q is proposed but instead leaves space in the binding cleft for ligand to enter and might bond with the oxygen at the ester group. The experiments conducted with triple mutant S209F W130H I179Q failed to beat HADDOCK score for DM, however its experimental results could still increase PET degradation. Results from surface charge may indicate an increase in stability and binding affinity for the protein.

**Conclusions:** Among models implemented, DM S209F W130H seems the best model studied regarding BHET or PET binding. Despite Protein Engineering is a complex process, computational tools might provide a way of studying binding sites of hypothetical proteins.

**Supplementary information:** Supplementary data is available in annexes.

## 1 Introduction

The origin of thermoplastic polyesters goes back to 1928 when Julian Hill was able to pull fibers from molten polymer^1^, but the work was dropped out in favor of nylon. However, PolyEthylene Terephthalate (PET) was only developed and patented in the year of 1941 in England by John Rex Whinfield, James Tennant Dickson and their employer the Calico Printers Association of Manchester^2^.

PET is a high molecular weight polymer composed of ester bond-linked terephthalate (TPA) and ethylene glycol^3^. Due to its durability and other favorable physical properties, PET became one of the most widely extensively utilized plastics being present in fibers for clothing, containers for liquids and foods, glass fiber among lots of other things. However, the large amount of PET that is being accumulated in ecosystems is of great environmental challenge. In 2016 alone, the annual production of PET was 56 million tons^4^.

In the same year a novel bacterium *Ideonella sakaiensis* 201-F6 was isolated which can utilize PET as an energy and carbon source^4^. This bacterium secretes an enzyme that from the polymer PET produces mono-(2-hydroxyethyl) terephthalic acid (MHET), TPA and bis-2(hydroxyethyl) TPA (BHET).

Petases or PET-hydrolyzing enzymes are biomolecules that decompose PET into its building blocks belonging to the Hydrolases family, which includes Esterases, Lipases, and Cutinases. Other Polyester-degrading enzymes were known since 1975 (α-chymotrypsin)^5^ and 1977 (Lipase)^6^ are two examples. Despite the enzyme found in *Ideonella sakaiensis* being highly homologous to several Cutinases, it exhibits 5.5 to 120-fold higher activity against PET hydrolysis over the other enzymes while presenting a lower activity on p-nitrophenol-linked aliphatic esters, the preferred substrates for Lipases and Cutinases^7^.

Petases could provide an eco-friendly solution to PET accumulation in the environment. PET-binding and hydrolysis mechanisms are known from previous works and given such important information there is the possibility of bioengineering these molecules in order to improve their functioning. On top of that there is also the possibility of improving protein function by changing other factors such as temperature, molecular weight and hydrophobicity. All of these possibilities require detailed studies of both protein structure and ligand binding mechanisms.

The aim of this article is to provide better insights into structure and binding site of PET Hydrolase, particularly the exploration of its catalytic mechanisms, analysis of the binding of ligands in models of hypothetical mutated structures and prediction of binding affinity of ligands using computational methods. More specifically, mutations were proposed that could lead to an increase in the catalytic activity of PET hydrolase.

This works contributes to the scrutiny of Molecular Dynamics (MD) behind the catalytic activity of PET hydrolase providing to the scientific community important information that can someday lead to a world without PET plastic in the environment.

### 1.1 Related Work

Given the importance of this subject as a current environmental challenge and possible improvement, there is a lot of related work already published. In 2017 Xu Han et al.^7^ published an article that serves as a base for the detailed mechanism of the catalytic activity of PETase. On top of that in 2018, Harry P. Austin et al. were then able to engineer a Double Mutant (DM) protein based on Cutinase protein family morphology, which highly increases PET degradation. Also, in 2018, Yuan Ma et al.^8^ published a study in which they proposed 3 mutations R61A, L88F, and I179F mutants, which were obtained with a rapid cell-free screening system and exhibited 1.4-fold, 2.1-fold, and 2.5-fold increases. These studies are the ones that will serve as a base for this project, however the research has not stopped since then.

In 2018 Hyeoncheol Francis Son et al.^9^ proposed the IsPETaseS121E/D186H/R280A variant, which was designed to have a stabilized β6-β7 connecting loop and extended subsite IIc, had a Tm value that was increased by 8.81 °C and PET degradation activity was enhanced by 14-fold at 40 °C.

More recently, in 2021 Andrew Renninson et al.^10^ have used sequence and structure-based bioinformatic tools to identify mutations to increase the thermal stability of the enzyme so as to increase PET degradation activity during extended hydrolysis reactions. Also, in 2021 Valentina P. et al.^11^ suggested a workflow for the improvement of bacterial PET Hydrolyzing enzymes and were able to produce a higher hydrolytic activity on PET.

### 1.1 Insights from Previous Work

Facts lead to the belief that PETase evolved from the Cutinase family since it presents a certain degree of sequence and structural homology^7^. In Figure 1 is shown the main changes that lead to such an increase in catalytic activity of the PET compounds. Mainly the appearance of disulfide bridge DS1 (C174-C210) which is inexistent in other Cutinases and the mutation of Histidine 184 into Serine 185. It seems this mutation creates space for different conformers of Tryptophan 156. Both of these mutations, as shall be discussed, were essential for increased catalytic activity related with PET.

**Figure 1.**
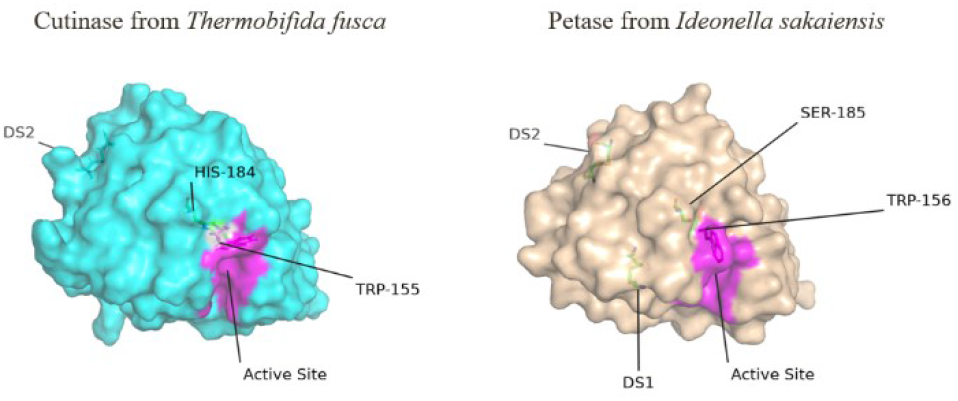
On the left side the 4CG2 Cutinase obtained from *Thermobifida fusca*. On the right side the 5XG0 from *Ideonella sakaiensis*.

To further understand the binding and catalytic mechanisms of this protein amino acids were highlighted according to their relation to catalysis as described in previous work^7^. Figure 2 shows the most import residues for binding and catalytic activity. The ligand is supposed to bind to A131, H208, W156, I179, W130, Y58, and M132, amino acids colored in magenta. The catalytic activity takes places by S131, H208, D177, present in protein surface and colored in cyan. While S131 seeks positive center and is at hydrogen-bond distance to be polarized by H208 which is in turn stabilized by D177. The backbone NH groups of M132 and Y58, colored in orange, constitute an oxyanion hole that holds ligand in place while being cut. The disulfide bridge DS1, specific of Petase, is essential for the function and is colored in Green. Finally, without background color, Tryptophan 156 is T-stacked to the benzene ring of ligand, facilitating rotation of reaction outputs after being cut due to its wobbling effect.

**Figure 2.**
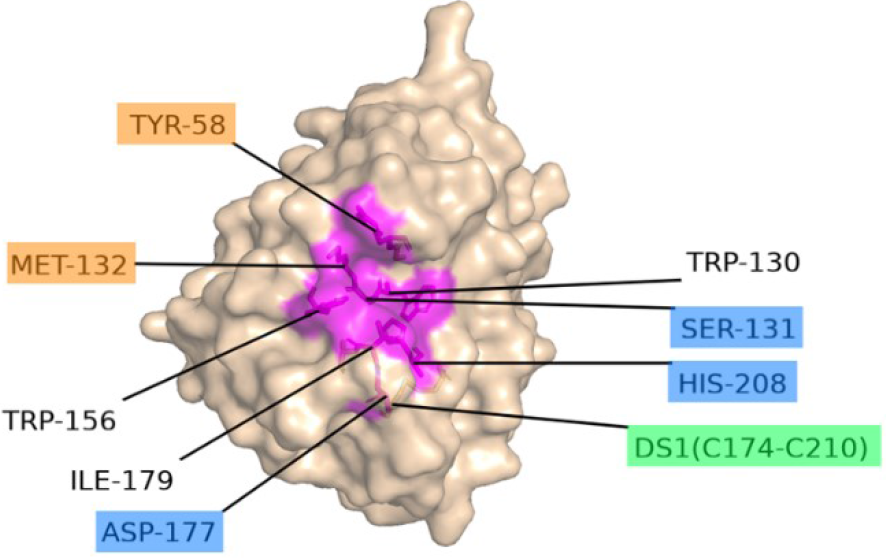
Chain A of 5XG0 PET hydrolase. The binding site is colored in magenta. The catalytic triad is colored in cyan. Oxyanion hole is colored in orange. Disulfide bridge is colored in green.

The same authors also tested a group of single mutations in order to understand how they would affect protein catalytic activity. In Figure 3 is possible to see the percentage of relative activity of protein byproducts for each mutant protein. As predicted, Serine 131 is essential for catalysis. Disrupting the disulfide bridge, also results in a nonfunctional protein. The remaining amino acids are essential for the binding of ligand but not for the catalysis. Lastly, disrupting the oxyanion hole on Tyrosine 58 and Threonine 59 mainly disturbed TPA production but not as many MHET. This indicates these amino acids are not essential for PET catalysis into MHET, but is required for MHET Catalysis into TPA.

**Figure 3.**
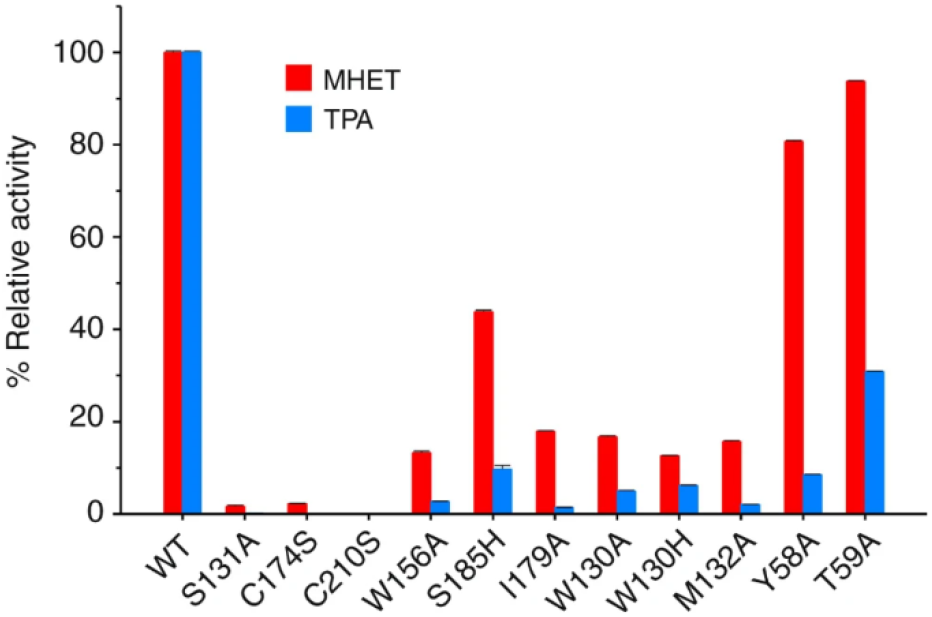
Production levels of MHET and TPA by each protein are presented as percentages of the Wild Type. Each measurement was conducted in triplicate, from which the average ± s.d. was calculated.

The Figure 2 was built and Figure 3 included in this work because it is crucial to understand ligand biding in order to improve catalytic activity. These amino acids are the ones the most affect the binding of ligand, and one should be very careful on their modification since that can have a drastic effect in the catalytic activity or even lead to compromise binding.

## 2 Material and Methods

In a first step the Wild Type (WT) structure was analyzed using PyMOL, a user-sponsored molecular visualization system, maintained and distributed by Schrödinger, from which multiple images and videos were rendered in order to elucidate about protein structure and ligand docking. Figure 4 shows the structure of the protein and its 3 chains A, B and C.

**Figure 4.**
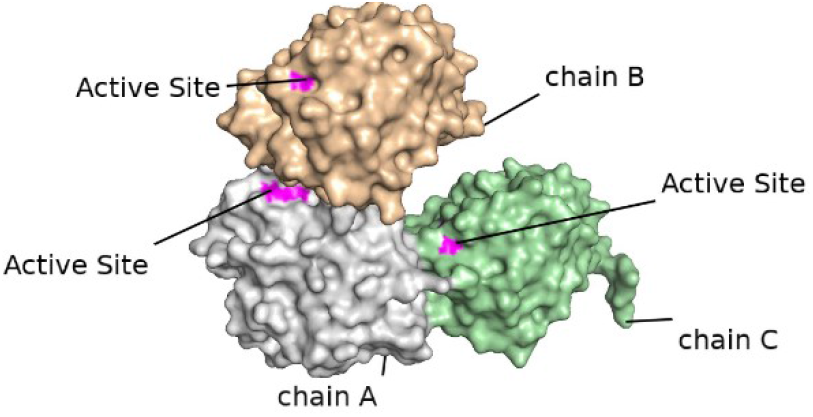
5XG0 - WT PET hydrolase from Ideonella sakaiensis.

Previous works were analyzed for insights regarding protein structure, function, the ligand binding site and catalytic activity from which were extracted a list of binding and active residues.

Based on residues identified and on the binding, activity described in previous work^7^, PyMOL was used to manually place a BHET ligand in the binding pocket. Then the proteomic strategies proposed by Rosário et al.^12^ were followed. The binding site was then analyzed and compared with results from HADDOCK (High Ambiguity Driven protein-protein DOCKing) an information-driven flexible docking approach for the modeling of biomolecular complexes which allows the study of molecular recognition and prediction of the binding mode and binding affinity of a complex formed by two or more constituent molecules with known structures.

Based on the work by Harry P. Austin et al.^13^ two improving mutations for the PET degrading protein were implemented based on the morphology of Cutinase from *Thermobifida fusca* which are known to increases its activity by around 5.5-fold in crystallinity change given a PET coupon. These mutations were S209F and W130H. This means that in position 209 Serine was mutated into Phenylalanine and in position 130 Tryptophan was mutated into Histidine.

Using Modeller, a software for homology or comparative modeling of protein three-dimensional structures^14,15^ 10 possible models for this Double Mutant (DM) protein were built, from which one was chosen that is most similar to both the WT Petase and Cutinase. In order to do that, the models were all aligned to the Wild Type Petase, the distribution of scores were normalized by the mean, and then the same was done for the Cutinase. To choose the best performing model, a sum of the normalized values was proposed as score and the model with the most negative value was chosen. This model basically narrows the binding cleft and opens space deeply into the active site.

The chosen model was used to simulate BHET binding with HADDOCK. Results were compared with WT.

Based on another work by Yuan Ma et al.^8^ another mutation, I179F, Isoleucine mutated into Phenylalanine, was implemented that increases PET degradation by 2.5-fold. 3D models were built for this single mutant using Modeller and the one with best DOPE score was selected.

The Mutant I179F model was then input into HADDOCK and compared with the best solution found for Wild Type. Despite showing great HADDOCK score, the W130 residue was identified to be obstructing the binding cleft, so a proposed solution was combining I179F with the DM S209F W130H, which opened space into the active site by mutating Tryptophan 130 into Histidine.

The models were once again generated and the binding evaluated using the same strategies. The solution don’t match the expectations due to the ligand seemingly having problems entering the active site. Suboptimal solutions on other HADDOCK clusters were evaluated, but results didn’t seem satisfactory. I179F that performed reasonably in the single mutant protein now seems to somehow be obstructing the way into the cleft that was narrowed by the double mutation. The model does not seem viable, so a proposed alternative was replacing this residue 179 into something less bulky and polar uncharged such as Glutamine, an amino acid that can also interact with Oxygen atoms present in the ligand.

Again, following the same strategy, models were built and the binding predicted for a single mutant I179Q and a Triple Mutant S209F, W130H, I179Q. The single mutant model doesn’t seem viable because there is not enough space for ligand binding due to the presence of W130, however in the Triple Mutant S209F, W130H, I179Q the ligand can now enter the catalytic site.

Note that BHET is a good predictor for a single unit, but at this point, the desired ligand was the PET polymer since some mutations introduced were supposed to make bonds to proximate units of the polymer. However, there were no PET polymer models available on PDB database so one was designed by combining four monomeric units using PyMOL builder.

Using HADDOCK, this new ligand was then tested for its binding to the known increased PET degrading activity I179, DM I209F W130H and the hypothetical triple mutant I209F W130H I179Q.

The surface charge distribution was also evaluated using the APBS plugin on PyMOL. The APBS is an implementation of the Poisson-Boltzmann equation, which describes the distribution of electric potential on a charged surface in contact with an ionic solution^16^.

Lastly, more information about the binding was needed and, to that end, a molecular dynamics simulation was created. These simulations are a popular technique for studying the atomistic behavior of molecular systems. Such simulations are complex and require a lot of setup steps. Reproduced below are the steps of Tutorial 5: Protein-Ligand Complex readily made available by “A Suite of Tutorials for the GROMACS-2018 Molecular Simulation Package” ^17^, which were the chosen methodology. The setup process for simulation requires the following steps:

- Preparation of the Topology for protein – which can be readily done using grep command from Unix/Linux and Gromacs;
- Preparation of the Topology for ligand – can’t be done simply by Gromacs so CHARMM General Force Field (CGenFF)^18,19^ was used, which performs atom typing and assignment of parameters and charges by analogy in a fully automated fashion^18,19^
- Building a complex with the protein and the ligand;
- Adding a simulation Box, Solvation and Ions to the system;
- Perform Energy minimization;
- Equilibration of the protein-ligand complex;
- Simulate the system using Molecular Dynamics;
- Analyzing Protein-Ligand Interactions and Ligand Dynamics.

After running the MD simulation, It should able to better understand the binding site, the catalytic activity on WT vs Double Mutant and from there try to implement improving mutations. Also, it would be possible to understand if the proposed Triple Mutant would be expected to have improved catalytic activity.

### 2.1 Materials

The PDB model from the Crystal structure of a novel PET hydrolase from *Ideonella sakaiensis* 201-F6 used as reference for the WT protein can be found in Protein Data Bank under the reference 5XG0 (https://www.rcsb.org/structure/5XG0). The structure was solved at 1.58 Å resolution with an asymmetric unit of three chains denoted A, B and C and adopts the standard α/β-hydrolase fold.

The model from the Cutinase, a thermostable polyethylene terephthalate degrading hydrolase from *Thermobifida fusca*, can be found in Protein Data Bank under the reference 4CG2 (https://www.rcsb.org/structure/4CG2). The structure is composed of a chain only with a resolution of 1.44 Å.

Models build using Modeller didn’t have particular restraints other than using the chain A of the WT model 5XG0.

HADDOCK is a complex simulation toolbox requiring many input parameters. Considering that the active site is known and how it is expected to operate, the active residues were always set as input for the toolbox. The residues 156, 59, 130, 58, 132 were selected for the WT and 158, 61, 132, 60, 134 for the generated models. One should note that despite being shifted 2 units these numbers represent the same amino acids, because in the WT the sequence starts in index -1. That doesn’t happen in the generated models.

Regarding HADDOCK simulations, the number of structures needed to be larger, since in pilot experiments the standard number was not large enough to provide consistent results. The following parameters were changed:

- Number of structures for rigid body docking: 3000;
- Number of structures for semi-flexible refinement: 1000;
- Number of structures for semi-flexible refinement: 500;
- Clustering Method: RMSD, Cuttoff 4.5.

## 3 Results and Discussion

### 3.1.1 Binding of BHET with the Wild Type Protein

HADDOCK was used for the prediction of binding between BHET and the WT protein. In Figure 5 it can be seen that the output came back similar to what was predicted based on given theoretical information. Distances in images can be hard to read, but movie1 for the manual placing and movie2 for the HADDOCK prediction can be found in Supplementary Information (SI) and might be useful. From the images, the catalytic S131 is at 5.0 Å from the ligand oxygen atom, while H208 is at 4.4 Å. Regarding oxyanion hole distances, M132 is the most proximate at 3.2 Å and T59 at 4.2 Å to oxygen atoms in the ligand. Interestingly the ligand position came back very similar to what was expected.

**Figure 5.**
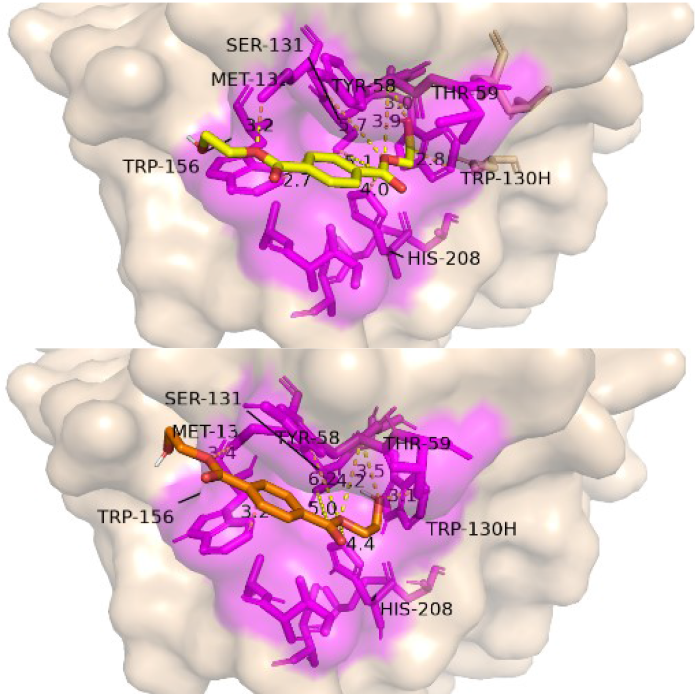
On top a representation of manual placement of BHET ligand near the binding site. On bottom the best HADDOCK solution for the binding of BHET with the WT protein.

### 3.1.2 Comparing Wild Type with Cutinase

The structure of the Wild Type protein was compared with the Cutinase. In Figure 6 is possible to see that in the Cutinase Phenylalanine 209 provides a narrower cleft for substrate distancing 5.1 Å, while in the Pet hydrolase the same distance is of 11 Å. In the work by Harry P. Austin et al.^13^ the authors also suggest this amino-acid provides new π-stacking and hydrophobic interactions to adjacent terephthalate moieties, but that can only be tested with PET polymer models and not BHET. On top of that, Histidine 129 allows the PET polymer to sit deeper within the active-site channel, maybe making it nearer to the catalytic S131. These results make sense according to what was reported in the previous work. In Table I is presented the results of aligning generated models with existing models for Wild Type (5XG0) and Cutinase (4CG2). Scale and centering of both alignment results were performed and M1 was chosen as the model most similar to both proteins, and picked to further testing.

**Figure 6.**
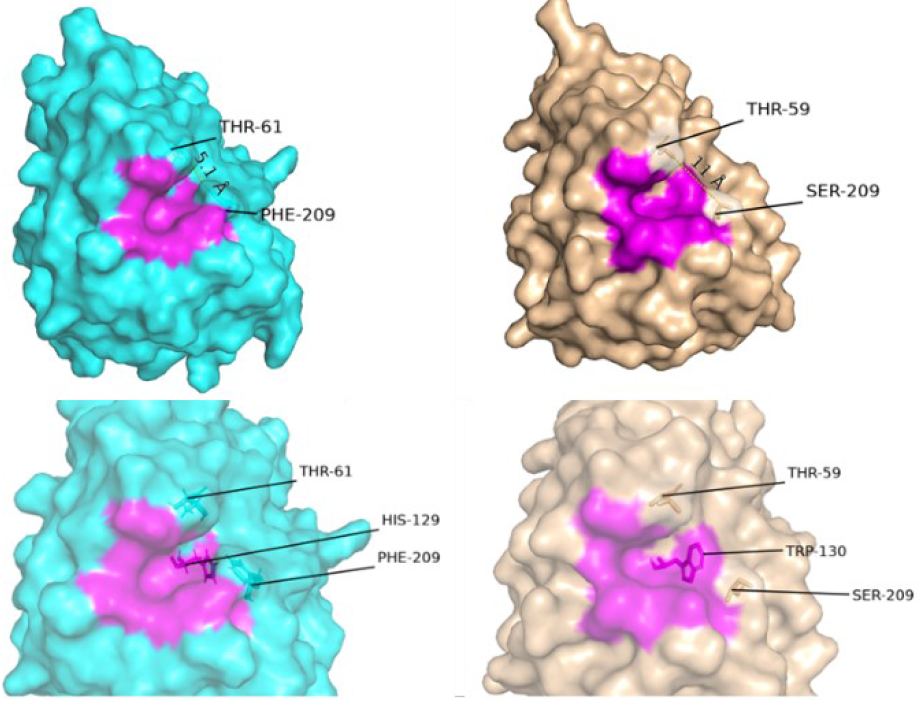
Comparison of Cutinase on Left, with Wild Type on Right. The mutated amino acids are also named in the figure.

**Table I.** Root Mean Square Deviation (RMSD) of the alignments of the generated models with the known structures.

| Model | DOPE score | Align WT<br>5xg0 | Align<br>4cg2 | Align 5xg0<br>(Norm) | Align 4cg2<br>(Norm) | SUM<br>(Norm) |
| --- | --- | --- | --- | --- | --- | --- |
| M1 | -27971,477 | 0,155 | 0,906 | -1,222 | -1,156 | -2,378 |
| M9 | -27829,781 | 0,157 | 0,927 | -0,970 | -0,260 | -1,230 |
| M8 | -27865,613 | 0,172 | 0,884 | 0,920 | -2,094 | -1,175 |
| M7 | -27716,043 | 0,166 | 0,917 | 0,164 | -0,687 | -0,523 |
| M5 | -27839,516 | 0,159 | 0,947 | -0,718 | 0,593 | -0,125 |
| M10 | -27836,992 | 0,164 | 0,942 | -0,088 | 0,380 | 0,291 |
| M4 | -27668,131 | 0,159 | 0,960 | -0,718 | 1,147 | 0,429 |
| M2 | -27863,047 | 0,169 | 0,936 | 0,542 | 0,124 | 0,665 |
| M3 | -27886,307 | 0,163 | 0,957 | -0,214 | 1,019 | 0,805 |
| M6 | -27927,299 | 0,183 | 0,955 | 2,305 | 0,934 | 3,239 |
| AVG.: |  | 0,165 | 0,933 |  |  |  |
| STD.DEV.: |  | 0,008 | 0,023 |  |  |  |

### 3.1.3 DM S209F W130H Binding of BHET

In Figure 7 the morphological changes that took place with the mutations from the WT to the DM protein are visible. It is important to refer the narrowing of the binding cleft by the Phenylalanine 209, as shown in blue; and the deepening of the binding site by the Histidine 130, as shown in wheat color.

**Figure 7.**
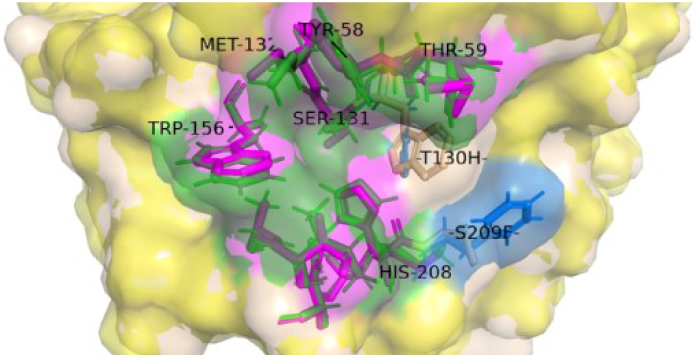
Morphological changes that took place in DM Petase. Residues in Magenta belong to Wild Type. Residues in Green belong to DM Model. Residues in Wheat color belong to Wild Type. Residues in Blue belong to DM Model.

The best HADDOCK solution for binding the DM S209F W130H with BHET is shown in Figure 8 (movie3 in SI). Here are some interesting changes. Firstly, it seems the ligand assumes a new position, this time being bound deeper into the pocket. It is translated to the right and flipped across the protein ligand axis. Because of this W156 is no longer near the benzene of BHET, but it is important to consider that in PET another unit can be connected to the left and W156 might stablish a π–π interaction with it. Something similar is supposed to happen to the right. The new mutated F208 might provide an aromatic interaction with next PET unit of the polymer. Catalytic S131 that was at 5.0 Å distance in Wild Type is now at 3.8 Å distance to the other side of the ligand and the cutting might take place there. Also new possible hydrogen bonds seem to appear at 3.2 Å and 3.0 Å with this placing of the ligand. This may also help explain why HADDOCK assigns a better score for this new placement of ligand.

**Figure 8.**
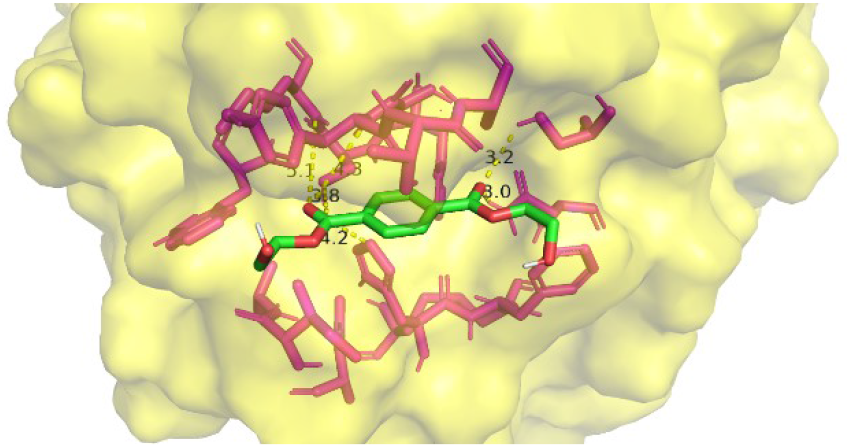
Best HADDOCK solution for binding of BHET with DM S209F W130H.

From previous work, it is known that DM S209F W130H has a much higher catalytic activity and in this way, in Table II it can be found that, as expected, the HADDOCK score, Van der Waals energy, Electrostatic energy and Desolvation energy for ligand binding was much lower than WT.

**Table II.** HADDOCK Predicted binding affinity for BHET.

| Metric | WT | DM S209F,W130H |
| --- | --- | --- |
| HADDOCK score | -25,2 +/- 0,2 | -34,4 +/- 0,3 |
| Van der Waals energy | -16,0 +/- 1,2 | -21,6 +/- 0,1 |
| Electrostatic energy | -2,5 +/- 3,5 | -18,3 +/- 1,4 |
| Desolvation energy | -9,0 +/- 0,6 | -11,1 +/- 0,2 |

### 3.1.3 I179F Binding of BHET

In Figure 9 (movie 4 in SI) is the best HADDOCK solution for another improving mutation reported in bibliography^8^, I179F. Despite achieving a better PET degradation activity, the HADDOCK simulation didn’t perform so well for the model generated. As can be seen, W156 seems to be obstructing the way into the cleft and thus S131 ends up being too distant (6.7 Å) from the related Carbon in the ligand.

**Figure 9.**
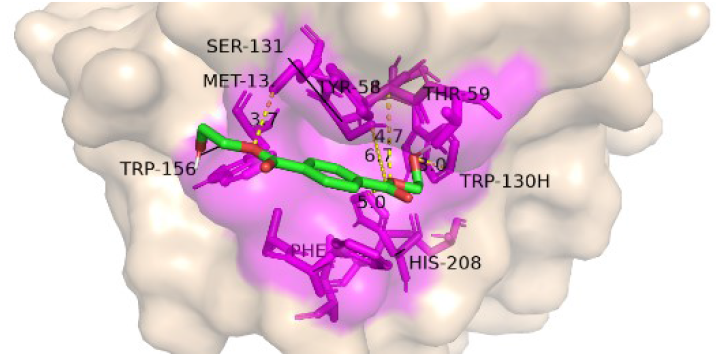
Best HADDOCK solution for binding of BHET with I179F.

In Table III it is shown that the HADDOCK score was way higher than WT, however Electrostatic energy was unexpectedly positive. These scores were expected to be better if PET were used as ligan instead of BHET because probably it fits better into the binding pocket.

**Table III.** HADDOCK Predicted binding affinity for BHET.

| Metric | WT | I179F |
| --- | --- | --- |
| HADDOCK score | -25,2 +/- 0,2 | -30.3 +/- 0.2 |
| Van der Waals energy | -16,0 +/- 1,2 | -15.9 +/- 0.2 |
| Electrostatic energy | -2,5 +/- 3,5 | 5.1 +/- 1.3 |
| Desolvation energy | -9,0 +/- 0,6 | -14.9 +/- 0.2 |

### 3.1.4 Could it be Possible to Combine both Solutions?

In an effort to improve the protein, these mutations were combined into a single model and thus a triple mutant S209F W130H I179F was built (Figure 10, movie5 in SI). However, this model doesn’t seem viable. The binding cleft that was already too narrow for single mutant I179F possibly became even more straight in the triple mutant. Suboptimal HADDOCK solutions were checked but they all showed similar problems. This way F179 seems to obstruct the way of ligand into cleft. Sometimes combining two good solutions can lead to a worse solution instead of making it better.

**Figure 10.**
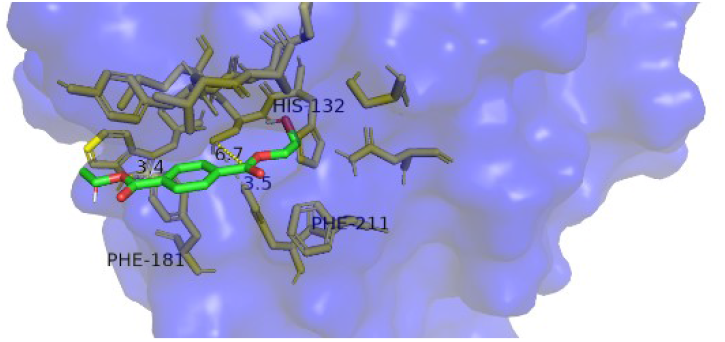
Best HADDOCK solution for binding of BHET with triple mutant S209F W130H I179F. The binding cleft seems to narrow for ligand binding and Catalytic S131 stays displaced at: a distance of 6.7 Å.

### 3.1.5 Proposing a Triple Mutant S209F W130H I179Q

The line of thought here was to replace residue 179 with Glutamine, since this amino acid is less bulky and polar uncharged, so it would interact with Oxygen atoms in ligand. In Figure 11 and Table IV the best HADDOCK Solution for single mutant I179Q are present. It seems, by the model and binding scores, that it doesn’t improve the WT catalytic activity. However, this mutation was also added to the DM S209F W130H, ending up with a triple mutant S209F W130H I179Q. In Figure 12 (movie5 in SI) is possible to see that the ligand can now enter into the catalytic site. Catalytic S131 is at 3.9 Å distance and oxyanion holes don’t seem disrupted. This solution seems to provide a working protein. The binding scores in Table IV indicate that it is not so great as DM, but one would need to experiment using a PET substrate to compare results with DM S209F W130H.

**Figure 11.**
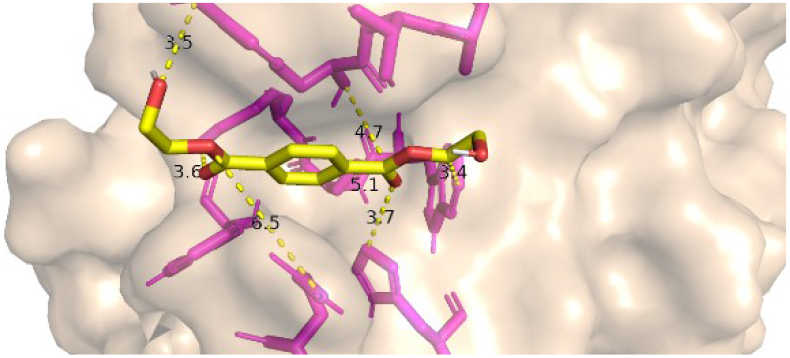
Best HADDOCK solution for binding of BHET with 179Q.

**Table IV.** HADDOCK Predicted binding affinity for BHET.

| Metric | I179Q | TM S209F W130H I179Q |
| --- | --- | --- |
| HADDOCK score | -25,6 +/- 0,1 | -32,6 +/- 0,1 |
| Van der Waals energy | -17,4 +/- 0,5 | -21,1 +/- 1,1 |
| Electrostatic energy | -10,3 +/- 3,8 | -10,5 +/- 3,2 |
| Desolvation energy | -7,1 +/- 0,2 | -10,5 +/- 0,7 |

**Figure 12.**
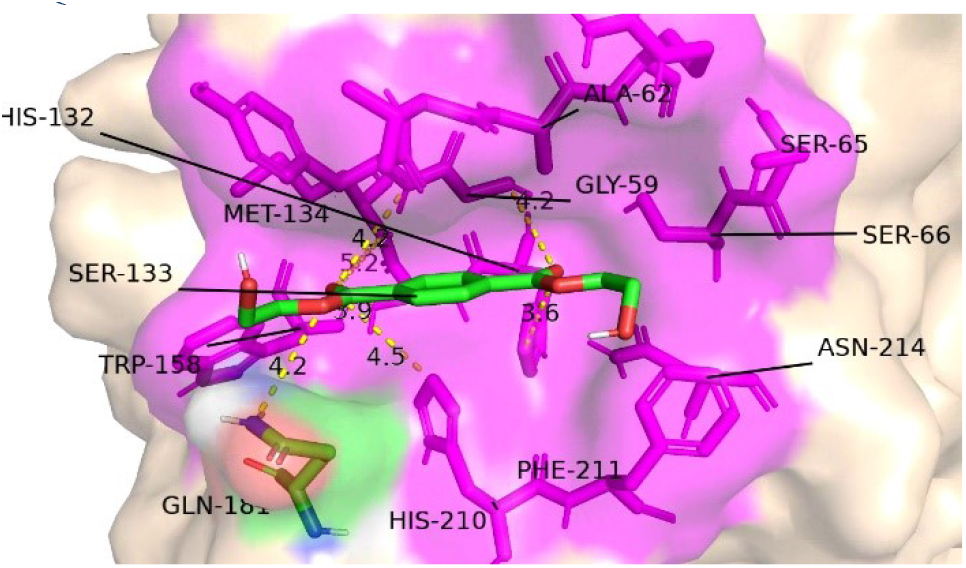
Best HADDOCK solution for binding of BHET with a triple mutant S209F W130H I179Q.

### 3.1.6 HADDOCK Scores for Binding PET

In Figure 13 a generated model for a chain of 2 monomers of PET is represented. In Table V are the HADDOCK binding scores of PET polymer with two units for proteins that seem functional from previous tests.

**Figure 13.**
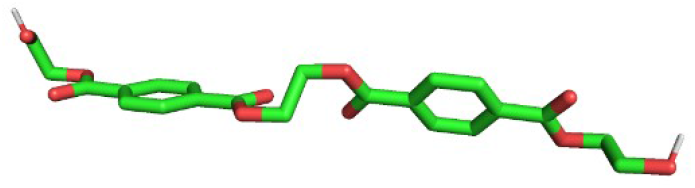
Model produced for PET ligand with 2 units chained.

**Table V.**
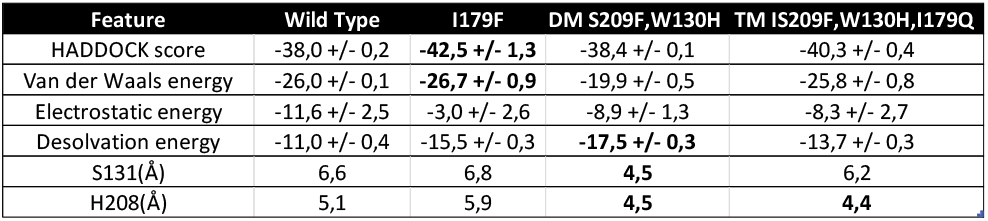
HADDOCK Predicted binding affinity for PET.

Regarding Table V. note that I179F has a -42.5 HADDOCK score while DM S209F W130H only has -38.4, however in DM S209F W130H catalytic Serine is at 4.5 Å, much nearer to the ligand. In fact, DM is known to be much more effective at degrading PET.

### 3.1.7 Surface Charge Distribution on PETase

The surface charge distribution was used as a tool to better understand the different conformations and binding preferences of the ligand. To that end, the Wild Type and DM S209F W130H were aligned and surfaces compared. It can be seen that with the mutation the groove is deepened, allowing the ligand to get closer to the catalytic site (Fig. 14).

**Figure 14.**
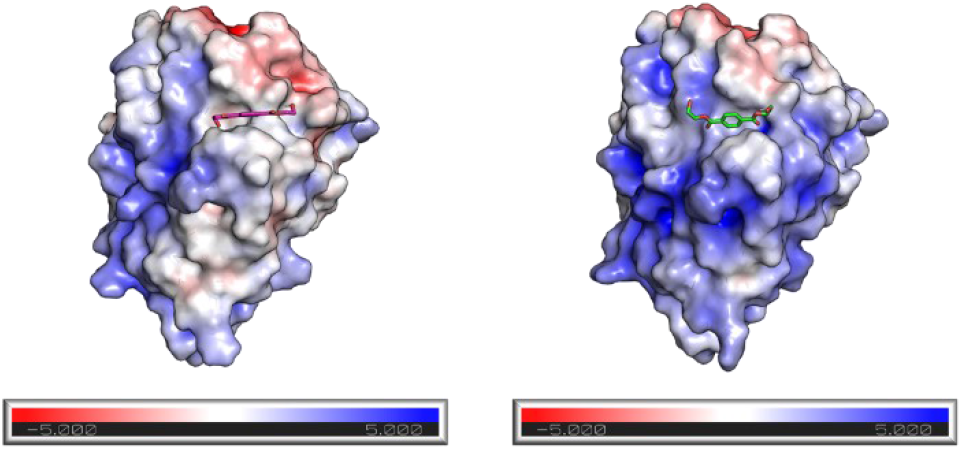
Surface Charge Distribution. On Left Wild Type. On Right Triple Mutant D83N D89N D157N.

**Figure 15.**
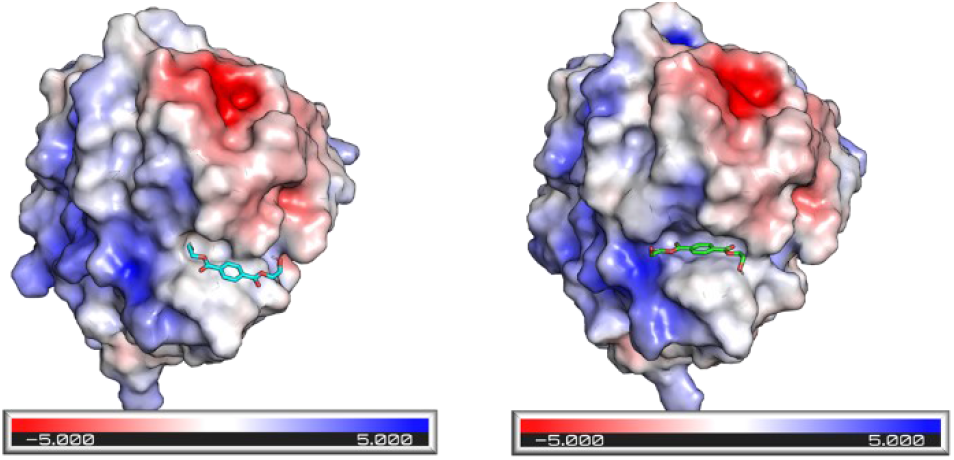
Surface Charge Distribution. On Left Wild Type. On Right DM S209F W130H.

However, it is also closer to the positive charges around the groove, and so, the molecule prefers a rotated conformation when compared to the WT, maybe due to the interaction with said charges.

Despite this change in orientation, the charges might help stabilize the molecule and increase the reaction rate. A mutant of the residues D83, D89 and D157 was suggested by Carola et al. (2021)^20^ to stabilize the active form of the protein where S131 becomes negatively charged. In the study they suggest that the residues cited be replaced by ones with positive side chain, which has a great impact on the surface charge of the protein and next to the groove.

### 3.1.8 Molecular Dynamics Simulation

Due to time constraints for this project, the MD simulation was only able to run with 3000 picoseconds, which was not enough for ligand to find the active site. A movie named gromacs.mp4 can also be found in SI which resumes the simulation. All simulation files are available in folder gromacs-simulation on SI. Figure 16 is one of the last frames of the simulation.

**Figure 16.**
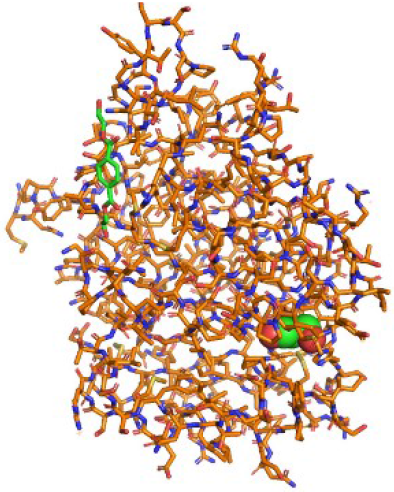
One of the last frames rendered with MD simulation of WT with BHET.

## 4 Conclusions

By using databases such as Protein Data Bank and tools such as Modeller, HADDOCK, PyMOL and Gromacs it is possible to assess binding sites and mechanisms of binding regarding existing and hypothetical proteins with ligands. In the context of this work, they were applied to the problem of bioengineering PETase.

It was possible to gain better insights regarding the binding mechanism of this protein and its mutations with BHET and extrapolate with what would happen with PET. However, it is important to stress that these methods are just estimations of what would happen and not definitive answers and do not preclude experimental assays.

Here is important to note that for instance I179F had a much better binding score than the DM, but it is known experimentally that it degrades PET 2.5-fold faster than WT while DM degrades PET at around 5-fold faster than WT. In this way, it is important to include other metrics such as distance of catalytic amino-acids to ligand in order to complement HADDOCK and free energy scores. Also, it seems by its morphology BHET might not be the best ligand for I179F. It is expected that these scores would fare better if using PET instead of BHET because the ligand should fit better into the binding pocket, and that was verified at a later phase of the project.

An improving mutation that would obviously outperform the Double Mutant S209F W130H was not found, but this study shows computationally that the Triple Mutant S209F W130H I179F and I179Q would not be good candidates since the cleft seems occluded to a good ligand fitting.

The experiments with triple mutant S209F W130H I179Q failed to beat HADDOCK score for DM, however it would be interesting to see if its experimental results would increase PET degradation somehow.

One should always be aware that protein engineering is a complex process and even in this project it was shown that combining two viable and improving mutations can lead to an inviable solution. Also changing just one residue can have unexpected consequences in structure and consequently on function.

In projects like this is important to know that the error is cumulative. There is error given by the degree of accuracy of the PDB model (resolution), then there is error regarding the prediction of 3D models, the error from HADDOCK simulations and so on. It is important to always work with most accurate results one can find and always find ways to reduce the amount of error in every step.

The results drawn from the surface charge rate were not analysed quantitatively, but they provide guidance to the understanding of the biochemistry underlying the process. Therefore, it can be inferred that the deepened grove on the DM might also work to stabilize the negative side chain on the catalytic site, leading to the observed increase in reaction rate.

Finally, we would like to remark that some details are hard to simulate and/or computationally demanding. For instance, the W156 wobbling effect is extremely hard to simulate and Gromacs simulations can take multiple days to run.

This project helps to clarify the binding mechanisms of PETase and provides information that may lead to an improvement in its catalytic activity.

## 5 Future Work

Regarding Pet hydrolase catalytic improvement there a lot more that can be done. Firstly, it would be interesting to achieve a better understanding of the binding and catalytic mechanisms of both WT and DM proteins before proposing more profound mutations. That could be done by finishing the Molecular Dynamics simulations and further analyze the respective data.Also, there are other ways of improving the protein such as making it more efficiently at other temperatures and pH that could be studied.

## 6 Limitations

Regarding improvement on the catalytic activity of PET hydrolase by protein engineering, it is important to refer that protein modeling, ligand binding simulations and molecular dynamics are complex processes requiring a lot of computational power. Also, in the context of this project limited time was available and thus it was not possible to deeply explore the binding mechanism and proposed more improvements. In order to achieve better and more accurate conclusions more time and access to more computational power would be needed.

## 7 Acknowledgments

The FP7 WeNMR (project# 261572), H2020 West-Life (project# 675858), the EOSC-hub (project# 777536) and the EGIACE (project# 101017567) European e-Infrastructure projects are acknowledged for the use of their web portals, which make use of the EGI infrastructure with the dedicated support of CESNETMCC, INFN-PADOVA-STACK, INFN-LNL-2, NCG-INGRIDPT, TW-NCHC, CESGA, IFCA-LCG2, UA-BITP, SURFsara and NIKHEF, and the additional support of the national GRID Initiatives of Belgium, France, Italy, Germany, the Netherlands, Poland, Portugal, Spain, UK, Taiwan and the US Open Science Grid.

